# Stabilizing the Closed SARS-CoV-2 Spike Trimer

**DOI:** 10.1101/2020.07.10.197814

**Authors:** Jarek Juraszek, Lucy Rutten, Sven Blokland, Pascale Bouchier, Richard Voorzaat, Tina Ritschel, Mark J.G. Bakkers, Ludovic L.R. Renault, Johannes P.M. Langedijk

**Author notes:** These authors contributed equally (Jarek Juraszek, Lucy Rutten).

## Abstract

The trimeric spike (S) protein of SARS-CoV-2 is the primary focus of most vaccine design and development efforts. Due to intrinsic instability typical of class I fusion proteins, S tends to prematurely refold to the post-fusion conformation, compromising immunogenic properties and prefusion trimer yields. To support ongoing vaccine development efforts, we report the structure-based design of soluble S trimers, with increased yields and stabilities, based on introduction of single point mutations and disulfide-bridges. We identify two regions in the S-protein critical for the protein’s stability: the heptad repeat region 1 of the S2 subunit and subunit domain 1 at the interface with S2. We combined a minimal selection of mostly interprotomeric mutations to create a stable S-closed variant with a 6.4-fold higher expression than the parental construct while no longer containing a heterologous trimerization domain. The cryo-EM structure reveals a correctly folded, predominantly closed pre-fusion conformation. Highly stable and well producing S protein and the increased understanding of S protein structure will support vaccine development and serological diagnostics.

## Introduction

Development of effective preventative interventions against the SARS-CoV-2 virus that causes the ongoing COVID-19 pandemic (Zhou, Yang et al. 2020, Zhu, Zhang et al. 2020) is urgently needed. The viral surface spike (S) protein is a key target for prophylactic measures as it is critical in the viral life cycle and the primary target of neutralizing antibodies (Brouwer, Caniels et al. 2020, Chen, Hotez et al. 2020, Liu, Wang et al. 2020, Yuan, Wu et al. 2020). S is a large, trimeric glycoprotein that mediates both binding to host cell receptors and fusion of virus and host cell membranes, facilitated by its S1 and S2 subunits respectively (Bosch, van der Zee et al. 2003, Li 2016, Wrapp, Wang et al. 2020). The S1 subunit comprises two distinct domains: an N-terminal domain (NTD) and a host cell receptor-binding domain (RBD). For infection, S requires proteolytic cleavage by a furin-like protease between the S1 and S2 subunits (S1/S2), and by TMPRSS2 at a conserved site directly preceding the fusion peptide (S2’) (Bestle, Heindl et al. 2020, Hoffmann, Kleine-Weber et al. 2020). In the pre-fusion state, the S-protein’s RBD domains alternate between open (‘up’) and closed (‘down’) conformations, transiently exposing the receptor’s binding site, in the ‘up’ conformation, which can bind to human angiotensin-converting enzyme 2 (ACE2). Like other class I fusion proteins, the SARS-CoV-2 S protein is intrinsically metastable as a consequence of its ability to undergo these conformational changes to drive fusion.

The prefusion conformation of S as present on the infectious particle contains the epitopes for neutralizing antibodies and thus holds most promise as vaccine immunogen (Brouwer, Caniels et al. 2020, Chen, Hotez et al. 2020, Hsieh, Goldsmith et al. 2020, Liu, Wang et al. 2020, Robbiani, Gaebler et al. 2020, Yuan, Wu et al. 2020). Prefusion stabilization typically increases the recombinant expression of viral glycoproteins which facilitates the production of protein for (subunit) vaccines and improves the immune response elicited by recombinant protein and viral vector vaccines (Graham, Gilman et al. 2019). In recent years, efforts have been made to stabilize various class I fusion proteins through structure-based design (for review see (Graham, Gilman et al. 2019)). A particularly successful approach to enhance prefusion stability was shown to be the stabilization of the so-called hinge loop preceding the central helix (CH) which has been applied to a range of viral fusion glycoproteins, (Sanders, Vesanen et al. 2002, Krarup, Truan et al. 2015, Battles, Mas et al. 2017, Hastie, Zandonatti et al. 2017, Rutten, Lai et al. 2018, Rutten, Gilman et al. 2020). Stabilization of the hinge loop of the S proteins of SARS-CoV and MERS-CoV has been achieved by mutation of two consecutive residues to proline in the S2 subunit between the central helix (CH) and heptad repeat 1 (HR1) (Pallesen, Wang et al. 2017, Kirchdoerfer, Wang et al. 2018) and this approach (2P) has successfully been applied into the SARS-CoV-2 S protein (Wrapp, Wang et al. 2020). However, the S protein carrying these substitutions with additional furin site mutations (S-2P) remains unstable and strategies have recently been described to improve stability (Henderson, Edwards et al. 2020, Hsieh, Goldsmith et al. 2020, McCallum, Walls et al. 2020, Xiong, Qu et al. 2020). Comparison of the structure of SARS-CoV-2 S-2P (Walls, Park et al. 2020, Wrapp, Wang et al. 2020) with that of native SARS-CoV-2 S (Cai, Zhang et al. 2020, Ke, Oton et al. 2020) shows that the former adopts a more open conformation with one or more of the RBDs in the ‘up’ conformation. Although neutralizing antibodies have been mapped to the RBD in the up as well as in the down state, the antibodies that bind the conserved epitopes on RBD in the down state were described to have the highest neutralizing potency (Brouwer, Caniels et al. 2020, Liu, Wang et al. 2020, Robbiani, Gaebler et al. 2020). Therefore, stabilizing the S in its closed (3 RBDs down) state and arresting the first step in the conformational change may result in an improved vaccine immunogen.

Using structure-based design, we found novel stabilizing mutations in both the S1 and S2 domains. Combining several of the novel mutations resulted in a highly stable S trimer, S-closed, with increased expression that remained stable in the absence of a heterologous trimerization domain that is typically required in soluble S designs (Walls, Tortorici et al. 2016, Pallesen, Wang et al. 2017, Walls, Park et al. 2020, Wrapp, Wang et al. 2020). Assessment of its antigenicity and high-resolution EM confirm that this trimer adopts a closed conformation.

## Results

To stabilize the S-protein in the closed pre-fusion state, we took a rational approach based on the structure of S (Wrapp, Wang et al. 2020). We searched for cavity filling substitutions, buried charges and possibilities for forming disulfides. We expanded our search for proline and glycine mutations, beyond the previously described hinge loop in the S-2P variant. Mutations were identified computationally with Rosetta’s mutant design (Kuhlman, Dantas et al. 2003) and Bioluminate cysteine bridge scanning (Salam, Adzhigirey et al. 2014) followed by visual inspection of molecular interactions. Selected mutations are presented in Fig. S1 and fall in two structural categories – SD1 head mutations N532P, T572I, D614N and S2 loop mutations A942P, T941G, T941P, S943G, A944P, A944G (loop α13α14) and A892P (loop α10α11). Disulfides F888C+G880C (DS1) and S884C+A893C (DS2) were identified in loop α10α11.

S ectodomain variants with mutations according to Fig. S1 were expressed as single chain by mutation of the furin site (ΔFurin) and addition of a C-terminal foldon (Fd) trimerization domain with (S-2P) or without the two previously described stabilizing prolines in the hinge loop (Wrapp, Wang et al. 2020). Supernatants of Expi293F cells transfected with plasmids encoding the S variants were tested for trimer content (Fig 1A) and for RBD exposure of the S protein by ACE2 binding (Fig. 1B). All mutations significantly increased trimer yields and ACE2 binding of the S protein for either the ΔFurin or S-2P variants. A strong effect was observed with T941P, A942P and A944P. A942P showed a ~11-fold increase in expression for ΔFurin, and ~3-fold for S-2P (Fig. 1A). For T941P, A944G and K986P shorter retention times were observed likely indicating opening of the trimers. T941P and A944G showed the highest ACE2 binding amongst the α13α14 loop mutations and K986P resulted in a ~10-fold higher ACE2 binding, whereas the trimer yield was only ~3-fold higher than ΔFurin. This indicates that the RBD domains are more exposed.

**Figure 1.**
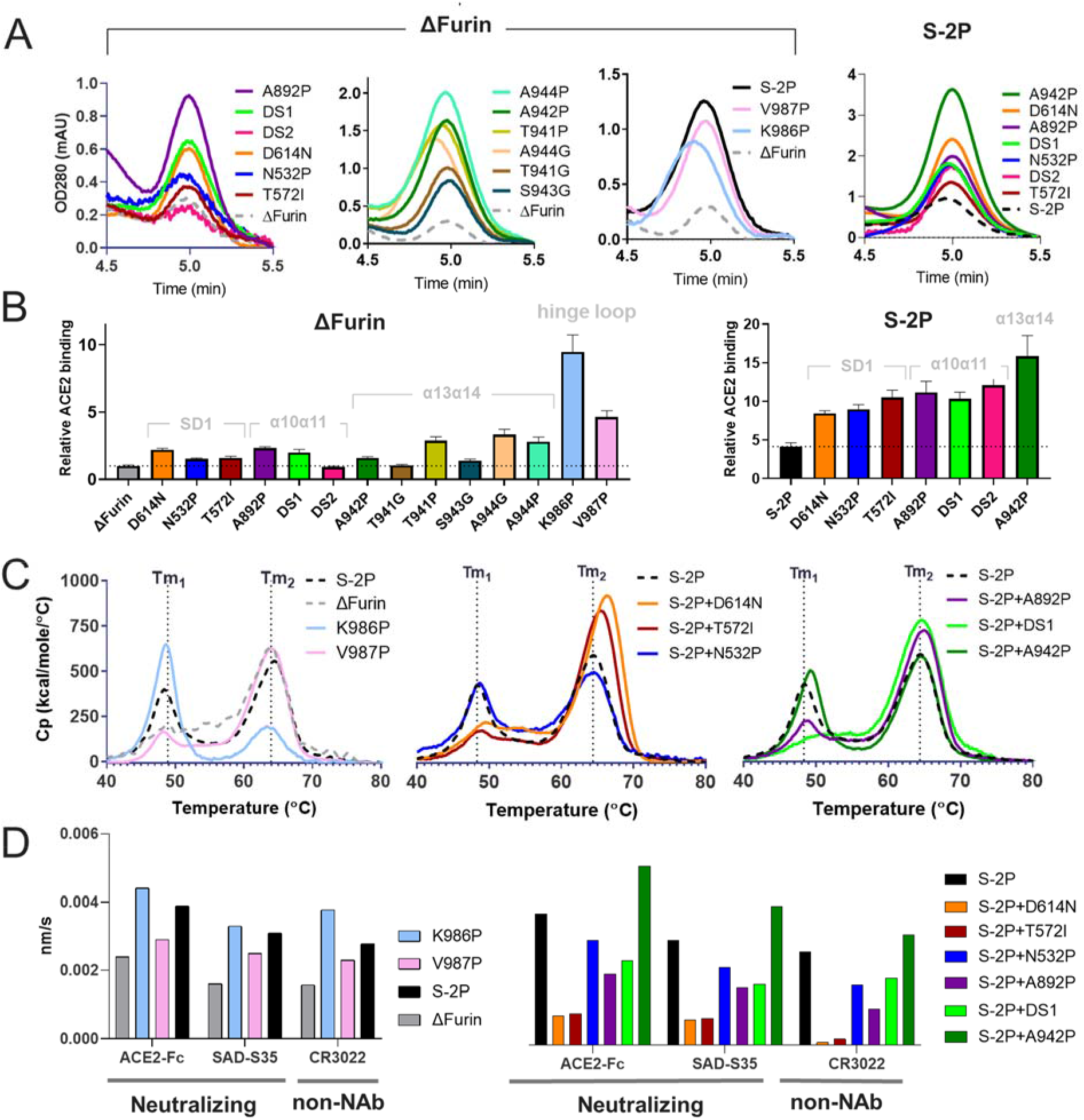
Characterization of SARS-CoV-2 S mutants containing single stabilizing mutations. (A) Analytical SEC of S variants showing the trimer peak (solid line) relative to the backbone (dashed line) (B). ACE2-Fc binding to S protein mutants based on AlphaLISA of ΔFurin variants (B, left panel) and S-2P (B, right panel). Data are represented as mean ± SEM. Mutants were grouped according to structural regions indicated in light grey. (C) Temperature stability of purified S trimers as measured by DSC. Two melting events are indicated by Tm1 and Tm2. (C, left panel) Uncleaved SARS2-S variants with furin site mutations (ΔFurin), with one stabilizing proline mutation in the hinge loop (ΔFurin K986P or ΔFurin V987P), and both proline mutations in the hinge loop (S-2P). (C, middle panel) ΔFurin variants with indicated mutations in S1 and (C, right panel) ΔFurin variants with indicated mutations in S2. (D) Binding of SAD-S35, ACE2 and CR3022 to purified S proteins measured with BioLayer Interferometry, showing the initial slope at the start of binding.

Stability of the single point mutants was further characterized with purified proteins using differential scanning calorimetry (DSC). First, we tested the contribution to stability of the individual proline mutations of the S-2P variant (Fig. 1C left panel, Table S1). All curves showed two melting events (T*m*_50_’s), albeit with different ratios. ΔFurin and additional V987P show a major T*m*_50_ at 64°C and a minor T*m*_50_ around 49°C. This is inverted for the K986P mutant in which the lower temperature transition is dominant. Combination of both prolines reduced the lower and increased the higher T*m*_50_. Amongst the S2 mutants (Fig. 1C central panel, Table S1) D614N diminished the peak height of the first T*m*_50_ and increased the second by almost 2°C, which was confirmed with differential scanning fluorimetry DSF (Fig. S2). A similar effect was observed for T572I and the loop stabilizing mutations A892P and DS1, albeit to a lesser extent (Fig. 1C right panel, Table S1). A942P which increased trimer yields hardly affected thermal stability in DSC (Fig. 1C right panel) nor DSF (Fig. S2).

RBD exposure was characterized by binding of ACE2; neutralizing antibody SAD-S35 and non-neutralizing antibody CR3022 that compete with ACE2. ACE2 and SAD-S35 can only bind RBD in the ‘up’ configuration and CR3022 can only bind when 2 RBDs are in the ‘up’ configuration (Yuan, Wu et al. 2020). The variant with K986P showed higher binding of SAD-S35, ACE2 and CR3022 than S-2P (Fig. 1D left panel), in accordance with the results obtained with SEC and AlphaLISA in supernatants. D614N and T572I showed very low binding to SAD-S35 and ACE2 and almost no CR3022 binding (Fig. 1D right panel), indicating more closed trimers. A892P improved trimer closure to a lesser extent, while A942P seemed to increase its opening. These mutants likely exhibit a mixture of closed, 1-up and 2-up structures.

The purified S variants with higher T*m*_50_ and lower Ab binding compared to S-2P also showed longer retention times in SEC (Fig. S3), in agreement with a more compact structure. The largest shift was caused by D614N. Since 614 corresponds to a position in the S protein subject to most extensive adaptation to the human host - D614G (Korber, Fischer et al. 2020), this variant was also tested for expression and stability. The results show that the D614G change has a very similar effect as D614N. Both increase trimer yields, increase Tm by ~2°C (Fig. S4) and reduce SAD-S35, ACE2 and CR3022 binding substantially (data not shown). Interestingly, both D614G and D614N full length variants show increased fusogenicity compared to wildtype D614 (Fig. S5). Next, we made variants with combined mutations to evaluate if the effects on expression, trimer closure and stability are additive. Being located in different protein domains, D614N and A892P were both selected for their positive effect on the stability of the spike and A942P for its strong increase in protein yields (Fig 1A). These mutations were combined with hinge loop prolines resulting in two combos, a quadruple mutant S-closed+Fd containing D614N+A892P+A942P+V987P and a quintuple mutant S-closed+Fd+K986P. Although both combos showed similar, approximately 5-fold improvement in yields compared to S-2P, the addition of K986P increased ACE2 binding (Fig. 2A). Interestingly, the quadruple mutant in which the foldon trimerization domain is deleted (S-closed), showed a 6.4-fold improvement in yields compared to S-2P. Its trimer peak was shifted towards longer retention times due to the smaller size which was confirmed by MALS analysis (Fig. 2A, Table S2).

**Figure 2.**
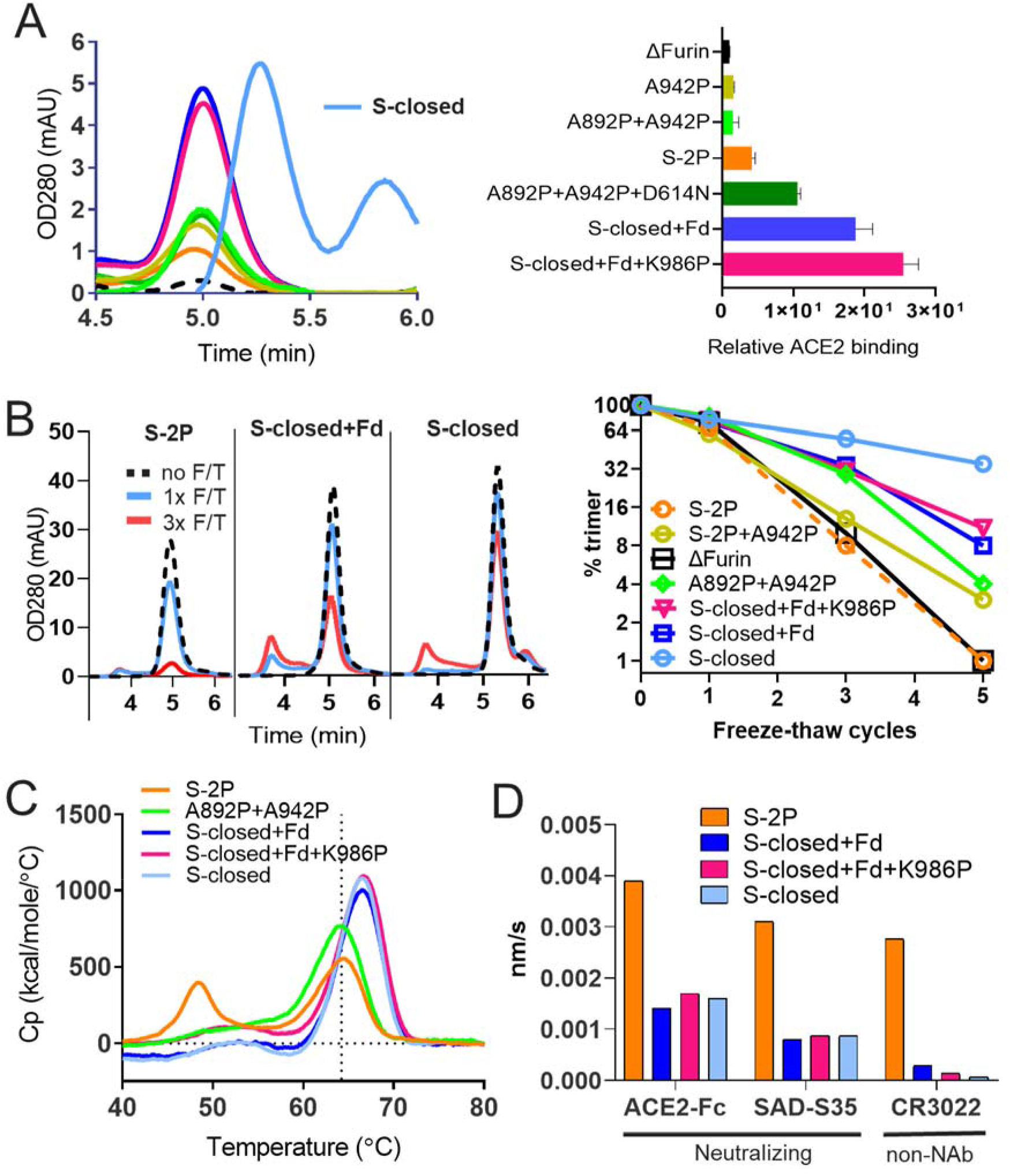
Characterization of SARS-CoV-2 S mutants containing combinations of stabilizing mutations. (A) Analysis of cell culture supernatant after transfection using analytical SEC (A, left panel) and ACE2-Fc binding based on AlphaLISA (A, right panel) of ΔFurin S with combinations of stabilizing mutations. Data are represented as mean ± SEM. (B) Freeze-thaw stability of purified uncleaved S trimers with indicated stabilizing mutations as measured by analytical SEC. Chromatograms are shown for non-frozen (dashed black line), 1 x frozen (light blue line) and 3 x frozen (red line). (C) Temperature stability of purified combined S trimer variants by DSC (D). Binding of SAD-S35, ACE2 and CR3022 to purified combined S trimer variants measured with BioLayer Interferometry, showing the initial slope at the start of binding.

S-closed with and without foldon were further characterized for resilience during repeated freeze-thaw cycles (Fig. 2B and Table S3). Analytical SEC showed that only 8% of the original S-2P trimer was present after 3, and only 1% after 5 cycles (Fig 2B, right panel). The trimer content is significantly improved for S-closed retaining 55% of intact trimers after 3, and 35% after 5 freeze-thaw cycles. Higher thermal stability of the combos was shown with only a single T*m*_50_ at about 66°C (Fig. 2C). All combos displayed decreased levels of ACE2 and antibody binding compared to S-2P control (Fig 2D, Fig. S6).

The S-closed+Fd quadruple mutant was then imaged by cryo-EM. A 2-steps 3D classification illustrates that out of 833,000 classified particles, ~80% was closed with all RBDs in the down state and 38% was categorized into a well-defined closed class and ~20% showed 1 RBD-up (Figure S7). Further processing of the 320,000 closed conformation particles allowed us to obtain a 3.06Å electron potential map (Figure S8). An atomic model that was built into the potential confirmed that S retains the prefusion spike conformation (Fig 3). The NTDs and RBDs density is less defined than for the rest of the map, suggesting flexibility in these regions. The closed structure (see Fig 3A) is highly reminiscent of the one previously solved by Walls et al (Walls, Park et al. 2020).

**Figure 3.**
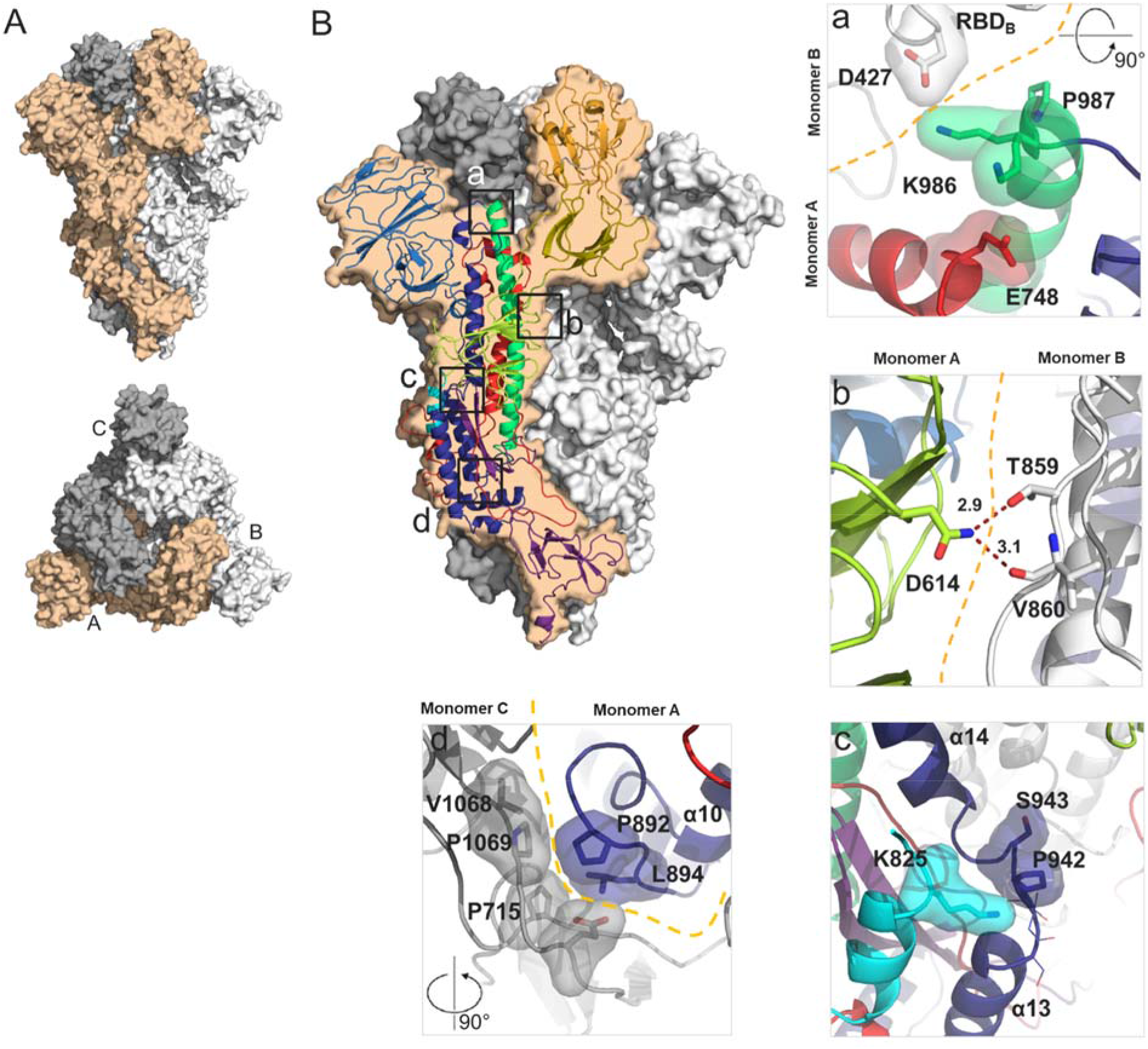
Structural characterization of S-closed. (A) Cryo-EM structure of the most abundant trimer class of S-closed-Fd – a closed S protein trimer. Monomers are colored in light orange, white and grey and the spike is shown from the side (upper panel) and from the top (lower panel) view. (B) Each of the four single point mutations introduced in S-closed-Fd shown in detail. Domains of the new structure are colored according to the same color code as used in Fig S1. Boundary between monomers has been additionally indicated with orange dashed line when applicable. Two possible rotamers of K986 are shown as one or the other could not be definitively assigned based on the density.

## Discussion

SARS-CoV-2 S proteins are unstable and although introduction of a double proline (K986P and V987P) in the hinge loop at the C-terminus of HR1 of S-2P was shown to improve stability of the prefusion conformation (Walls, Park et al. 2020, Wrapp, Wang et al. 2020), the S protein still suffered from instability (Fig 1C, 2B). We show that while each proline mutation increases trimer expression, the K986P variant also reduces the interaction with the S1 head releasing the RBD to the ‘up’ configuration based on increased ACE2 and CR3022 binding (Fig 1D) and a leftward shift in the SEC profile (Fig. 1A). The effect is partially compensated by the introduction of the second stabilizing mutation - V987P. We describe here two groups of substitutions that further stabilized the S-2P prefusion trimers and reduced RBD exposure. The first group of additionally novel stabilizing mutations was identified in the S2 HR1 region that undergoes an extensive conformational change during fusion. Stabilizing proline and glycine mutations in loops α10α11 and α13α14, and disulfides in loop α10α11 and between the central helix and C-terminus of HR1 showed significantly increased trimer yields. It is likely that the mutations in S2 facilitate folding of the protein during expression, by fixing loop conformations with otherwise intrinsic alpha helical propensity, necessary for driving the fusion conformational change. This strategy greatly improved prefusion S protein expression levels (Fig 1A). Recently, also Hsieh et al. in parallel to our study, demonstrated the stabilizing effect of the A942P and A892P mutations (Hsieh, Goldsmith et al. 2020). Mutations identified in the head region clustered near the SD1 subdomain. Interestingly, two of these mutations (D614N and T572I), aside of increasing expression in the S-2P variant, displayed a striking improvement of thermal stability (Fig. 1C) and exhibited very low RBD exposure (Fig.1D).

Based on the experimental characterization of single point mutants and their subdomain placement we selected four substitutions with low surface exposure (D614N, A892P, A942P and V987P) to create S-closed which doesn’t contain a heterologous trimerization domain and exhibits 6.4-fold increase in expression (Fig 2A), high thermal and freeze-thaw stability (Fig. 2B, C) and antigenicity reminiscent of a closed trimer (Fig. 2D). Both D614G and D614N variants show increased fusogenicity and stability (Fig. 1C, Fig. S5) which may be explained by a decrease in premature shedding of S1 (Zhang, Jackson et al. 2020). The interaction of D614 with the fusion peptide proximal region (FPPR) may play a role in stabilizing the spike during expression, as recently elucidated by a low pH structure (Zhou, Tsybovsky et al. 2020), in which D614 is in direct proximity of D839 and forms a hydrogen bond with the carbonyl of V860, suggesting aspartate protonation. In our new structure N614 forms two interprotomeric hydrogen bonds with the sidechain of T859 and backbone carbonyl of V860 (Fig. 3B), effectively mimicking the stabilizing effect of the protonated aspartate residue, without the pH dependence. It remains unknown why the D614G has a similar positive effect on stability, but perhaps the high cluster of negative charges in the interface between S1 and S2 destabilizes the protein and the 614N or 614G mutation reduces this repulsion. None of the stabilizing proline mutations modify the local structure of the protein, while A892P adds new interprotomeric interactions with the neighboring hybrid sheet, composed of both S1 and S2 strands (see Fig 3B). The stabilizing mutations and presence of K986 which interacts with Asp427 in the RBD of the neighboring monomer and Glu748 in S2 (Fig 3B) allow the S trimer to maintain predominantly the closed prefusion configuration.

The viral spike is mostly closed (Ke, Oton et al. 2020) and although many neutralizing antibodies are directed against RBD, the antibodies with the highest neutralizing activity bind RBD in the down state (Brouwer, Caniels et al. 2020, Liu, Wang et al. 2020, Robbiani, Gaebler et al. 2020). Similar to other class I fusion proteins, a closed conformation may be more reminiscent of transmitted virus and antibodies that recognize the closed state may be more important for protection as has been shown for HIV (Sanders and Moore 2017). Furthermore, the closed conformation can potentially induce antibodies with higher cross reactivity since the down state surface is more conserved. A stable closed S trimer with minimal non-exposed mutations and without a foldon that shows a significant increase in expression levels may advance development of novel (subunit) vaccine immunogens and further improve genetic vaccines, diagnostics or isolation of antibodies.

## Acknowledgements

This work benefited from access to the Netherlands Centre for Electron Nanoscopy (NeCEN) at Leiden University, an Instruct-ERIC center with assistance from Wen Yang and Frederic Bonnet. We thank Wouter Koudstaal for advice and assistance. We thank Ilona Bisschop, Martijn de Man, Ava Sadi, Anne-Marie de Gooyert, Lam Le and Annemart Koornneef for technical support.

## Author contribution

J.J., L.R. M.B. and J.P.L. designed the study, J.J. performed structure based design of mutations, T.R., S.B., P.B. and R.V. planned and / or performed biochemical assays and purifications, L.L.R.R. performed EM sample preparation, data collection, data processing and analysis, J.J., L.R., M.B., L.L.R.R. and J.P.L wrote the paper

## Methods

### Protein expression and purification

A plasmid corresponding to the semi-stabilized SARS-CoV2 S-2P protein (Wrapp, Wang et al. 2020) was synthesized and codon-optimized at GenScript (Piscataway, NJ 08854). A variant with a HIS tag and a variant with a C-tag were purified. The constructs were cloned into pCDNA2004 or generated by standard methods widely known within the field involving site-directed mutagenesis and PCR and sequenced. The expression platform used was the Expi293F cells. The cells were transiently transfected using ExpiFectamine (Life Technologies) according to the manufacturer’s instructions and cultured for 6 days at 370oC and 10% CO2. The culture supernatant was harvested and spun for 5 minutes at 300 g to remove cells and cellular debris. The spun supernatant was subsequently sterile filtered using a 0.22 μm vacuum filter and stored at 4°C until use. His – tagged SARS-CoV2 S trimers were purified using a two-step purification protocol by 1 mL or 5mL cOmplete His-tag columns (Roche). Proteins were further purified by size-exclusion chromatography using a HiLoad Superdex 200 16/600column (GE Healthcare).

## Antibodies and reagents

SAD-S35 was purchased at Acro Biosystems. ACE2-Fc was made according to Liu et al. 2018. Kidney international. For CR3022 the heavy and light chain were cloned into a single IgG1 expression vector to express a fully human IgG1 antibody. CR3022 was made by transfecting the IgG1 expression construct using the ExpiFectamine™ 293 Transfection Kit (ThermoFisher) in Expi293F (ThermoFisher) cells according to the manufacturer specifications. CR3022 was purified from serum-free culture supernatants using mAb Select SuRe resin (GE Healthcare) followed by rapid desalting using a HiPrep 26/10 Desalting column (GE Healthcare). The final formulation buffer was 20 mM NaAc, 75 mM NaCl, 5% Sucrose pH 5.5.

### Differential scanning calorimetry (DSC)

Melting temperatures for S protein variants were determined using a PEAQ-DSC system. 325 μL of 0.3 mg/mL protein sample was used per measurement. The measurement was performed with a start temperature of 20°C and a final temperature of 100°C. The scan rate 100°C/h and the feedback mode; Low (= signal amplification).

### Differential scanning fluorometry (DSF)

0.2 mg of purified protein in 50 μl PBS pH 7.4 (Gibco) was mixed with 15 μl of 20 times diluted SYPRO orange fluorescent dye (5000 x stock, Invitrogen S6650) in a 96-well optical qPCR plate. A negative control sample containing the dye only was used for reference subtraction. The measurement was performed in a qPCR instrument (Applied Biosystems ViiA 7) using a temperature ramp from 25–95 °C with a rate of 0.015 °C per second. Data were collected continuously. The negative first derivative was plotted as a function of temperature. The melting temperature corresponds to the lowest point in the curve.

### BioLayer Interferometry (BLI)

A solution of SAD-S35 at a concentration of 1 μg/mL and ACE2-Fc and CR3022 at a concentration of 10 μg/ml was used to immobilize the ligand on anti-hIgG (AHC) sensors (FortéBio cat#18-5060) in 1x kinetics buffer (FortéBio cat#18-1092) in 96-well black flat bottom polypylene microplates (FortéBio cat#3694). The experiment was performed on an Octet RED384 instrument (Pall-FortéBio) at 30 °C with a shaking speed of 1,000 rpm. Activation was 60 s, immobilization of antibodies 900 s, followed by washing for 600 s and then binding the S proteins for 300 s. The data analysis was performed using the FortéBio Data Analysis 8.1 software (FortéBio). Binding levels were plotted as nm shifts at 600s after S protein binding.

### Cryo-EM

#### Cryo-EM Grid Preparation and Data Collection

SARS-CoV-2 S protein samples were prepared in 20mM Tris, 150mM NaCl, pH7 buffer at a concentration of 0.15 mg/mL and applied to glow discharged Quantifoil R2/2 200 mesh grids before being double side blotted for 3 seconds in a Vitrobot Mark IV (Thermo Fisher Scientific and plunge frozen into liquid ethane cooled. Grids were loaded into a Titan Krios electron microscope (Thermo Fisher Scientific) operated at 300kV, equipped with a Gatan K3 BioQuantum direct electron detector. A total of 9,760 movies were collected over two microscopy sessions at The Netherlands Centre for Electron Nanoscopy (NeCEN). Detailed data acquisition parameters are summarized in Supplementary Table 4.

### Cryo-EM image processing

Collected movies were imported into RELION-3.1-beta (Zivanov, Nakane et al. 2018) and subjected to beam induced drift correction using MotionCor2 (Zheng, Palovcak et al. 2017) and CTF estimation by CTFFIND-4.1.18 (Rohou and Grigorieff 2015). Detailed steps of the image processing workflow are illustrated in Figure S8. Final reconstructions were sharpened and locally filtered in RELION post-processing (Fig. S8).

### Model building and refinement

The SARS-CoV-2 S PDBID 6VXX and 6VSB structures (Walls, Park et al. 2020, Wrapp, Wang et al. 2020) were used as starting models. PHENIX-1.18.261 (Liebschner, Afonine et al. 2019), Coot (Emsley, Lohkamp et al. 2010) and the Namdinator webserver (Kidmose, Juhl et al. 2019) were iteratively used to build atomic models. Geometry and statistics are given in Table S4. Final maps were displayed using UCSF ChimeraX (Goddard, Huang et al. 2018).

### AlphaLISA

Crude supernatants were diluted 300 times in AlphaLISA buffer (PBS + 0.05% Tween-20 + 0.5 mg/mL BSA). Then, 10 μL of each dilution were transferred to a 96-well plate and mixed with 40 μL acceptor beads, donor beads and ACE2-Fc. The donor beads were conjugated to ProtA (Cat#: AS102M, Perkin Elmer), which binds to ACE2Fc. The acceptor beads were conjugated to an anti-His antibody (Cat#: AL128M, Perkin Elmer), which binds to the His-tag of the construct. The mixture of the supernatant containing the expressed S protein, the ACE-2-Fc, donor beads, and acceptor beads was incubated at room temperature for 2 hours without shaking. Subsequently, the chemiluminescent signal was measured with an Ensight plate reader instrument (Perkin Elmer). The average background signal attributed to mock transfected cells was subtracted from the AlphaLISA counts. Subsequently, the whole data set was divided by signal measured for the SARS CoV-2 S protein having the S backbone sequence signal to normalize the signal for each of the S variants tested to the backbone.

### Analytical SEC

An ultra high-performance liquid chromatography system (Vanquish, Thermo Scientific) and μDAWN TREOS instrument (Wyatt) coupled to an Optilab μT-rEX Refractive Index Detector (Wyatt), in combination with an in-line Nanostar DLS reader (Wyatt), was used for performing the analytical SEC experiment. The cleared crude cell culture supernatants were applied to a SRT-10C SEC-500 15 cm column, (Sepax Cat# 235500-4615) with the corresponding guard column (Sepax) equilibrated in running buffer (150 mM sodium phosphate, 50 mM NaCl, pH 7.0) at 0.35 mL/min. When analyzing supernatant samples, μMALS detectors were offline and analytical SEC data was analyzed using Chromeleon 7.2.8.0 software package. The signal of supernatants of non-transfected cells was subtracted from the signal of supernatants of S transfected cells. When purified proteins were analyzed using SEC-MALS, μmMALS detectors were inline and data was analyzed using Astra 7.3 software package. For the protein component, a dn/dc (mL/g) value of 0.1850 was used and for the glycan component a value of 0.1410. Molecular weights were calculated using the RI detector as [C] source and mass recoveries using UV as [C] source.

### Cell-cell fusion assay

Quantitative cell-cell fusion assays were performed to ascertain the relative fusogenicity of the different D614 S protein variants by using the NanoBiT complementation system (Promega). Donor HEK293 cells were transfected with full-length S and the 11S subunit in 96-well white flat bottom TC-treated microtest assay plates. Acceptor HEK293 cells were transfected in 6-well plates (Corning) with ACE2, TMPRSS2 and the PEP86 subunit, or just the PEP86 subunit (‘Mock’) as negative control. All proteins were expressed from pcDNA2004 plasmids using Trans-IT transfection reagent according to the manufacturer’s instructions. 18hr after transfection, the acceptor cells were released by 0.1% trypsin/EDTA and added to the donor cells at a 1:1 ratio for 4 hr. Luciferase complementation was measured by incubating with Nano-Glo® Live Cell Reagent for 3 min, followed by read-out on an Ensight plate reader (PerkinElmer).

