## Supplementary Figures and Tables for "Stabilizing the Closed SARS-CoV-2 Spike Trimer"

### Supplementary Information

Jarek Juraszek[1]*, Lucy Rutten[1]*, Sven Blokland[1], Pascale Bouchier[1], Richard Voorzaat[1], Tina Ritschel[1], Mark J.G. Bakkers[1], Ludovic L.R. Renault[2], Johannes P.M. Langedijk#[1]

[1]Janssen Vaccines & Prevention BV, Leiden, the Netherlands

[2] NeCEN, Leiden University, Einsteinweg 55, Leiden, the Netherlands

*These authors contributed equally

#### Supplementary Figures

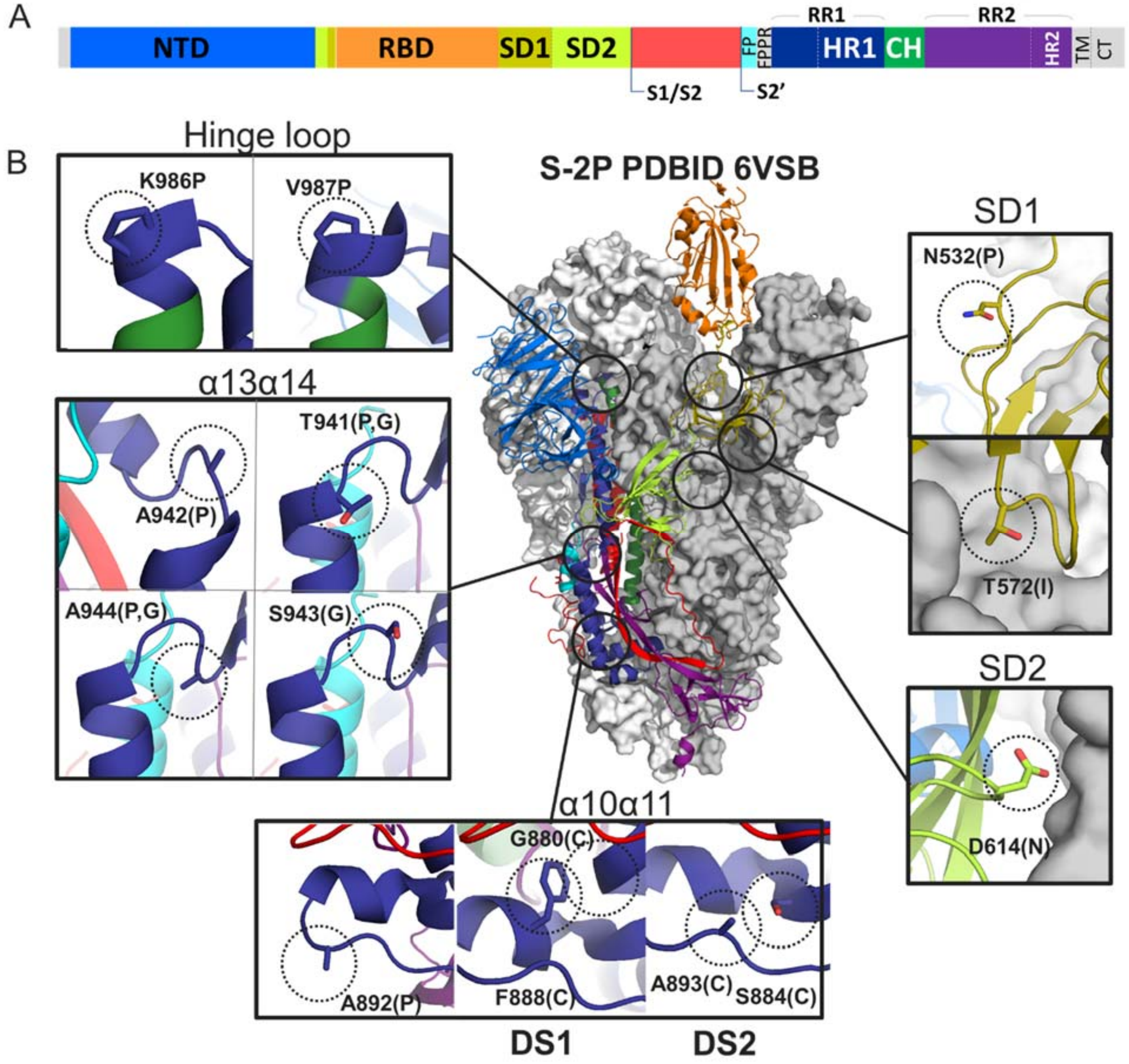


**Figure S1. Structure-based design of SARS-CoV-2 S stabilizing mutations**. (A) Domains of the SARS-CoV-2 S protein with indicated cleavage sites S1/S2 and S2'. (B). Stabilizing mutations identified in the S protein grouped by structural domains. Structure of S (PDBID 6VSB) colored according to panel A for one monomer. The other two monomers' surface is plotted in grey and white. K986P and V987P, the hinge prolines stabilizing the S-2P variant are also shown. Mutations applied to the indicated amino acids are indicated between brackets.

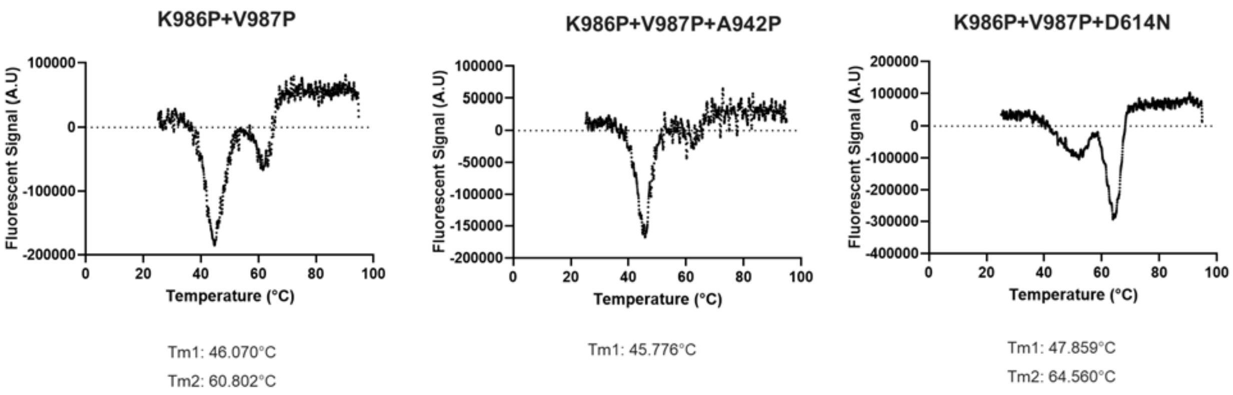


**Figure S2**. Differential scanning fluorimetry. Analysis of melting temperature (Tm) using differential scanning fluorimetry of purified S protein variants. The first order derivatives are plotted. Duplicate runs were averaged, and Tm determined as the lowest derivative value representing the $Tm_{50}$ value.

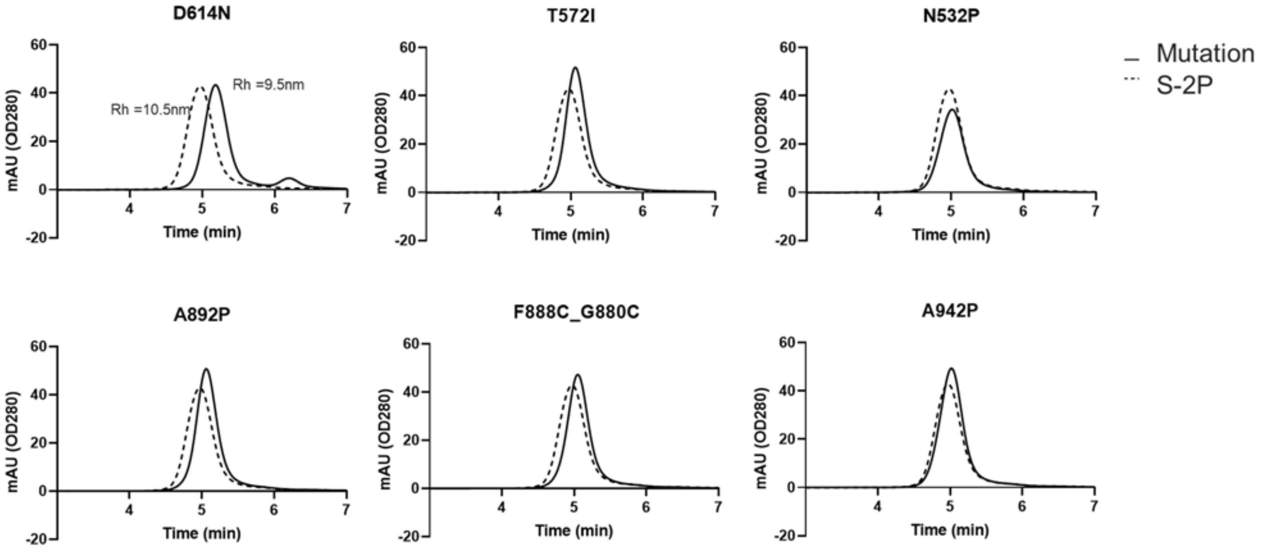


**Figure S3. Size exclusion chromatography of purified proteins**. Six SEC profiles of single mutation or disulfide variants of S-2P are compared with S-2P (dotted line).

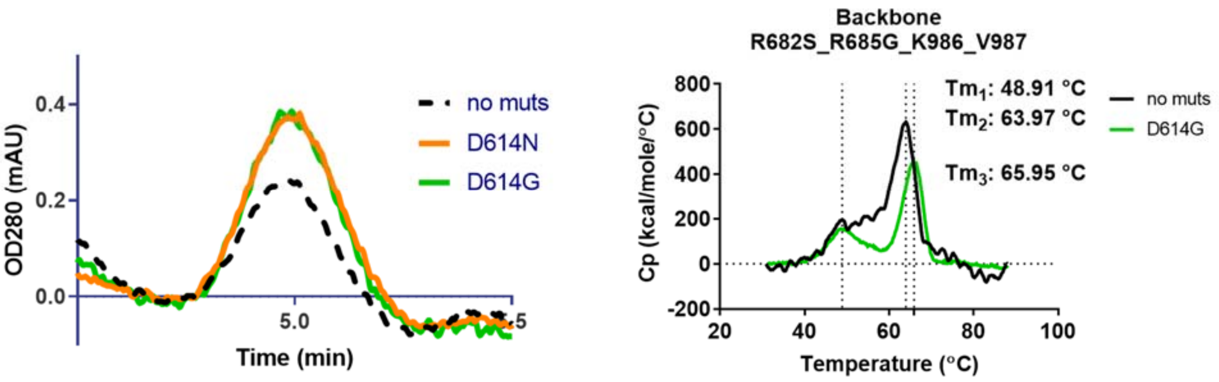


**Figure S4. SEC and DSC analysis of D614N/G mutants. (**A) Analytical SEC showing the trimer peak (solid line) relative to the backbone (dashed line). (B) Temperature stability of D614G variant trimer and the backbone trimer as measured by DSC. Two melting events are indicated by Tm1 and Tm2 for the backbone and Tm3 is indicated for the D614G variant.

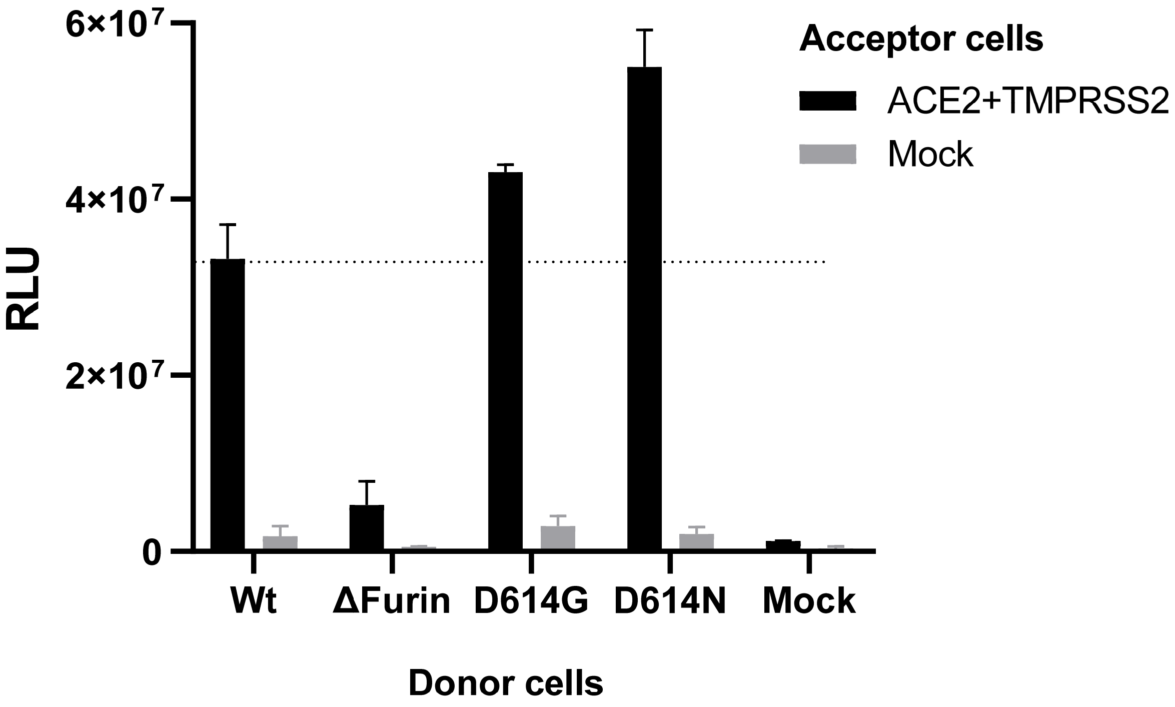


**Figure S5. Increased fusogenicity of D614N and D614G S variants.**

Quantitative cell-cell fusion assay in HEK293 cells using a split-luciferase approach. The level of fusion measured for wildtype S is indicated with a dotted line.

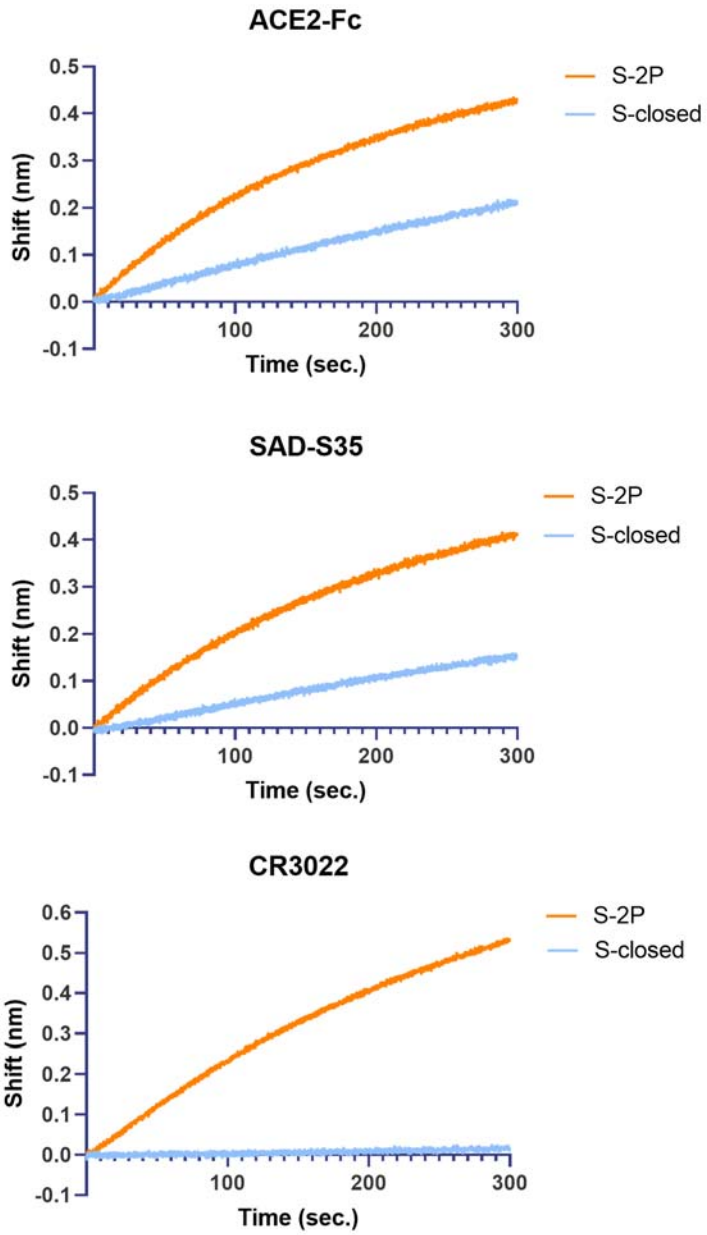


**Figure S6 BioLayer Interferometry using Octet.** Bio-Layer Interferometry curves for S-2P and S-closed

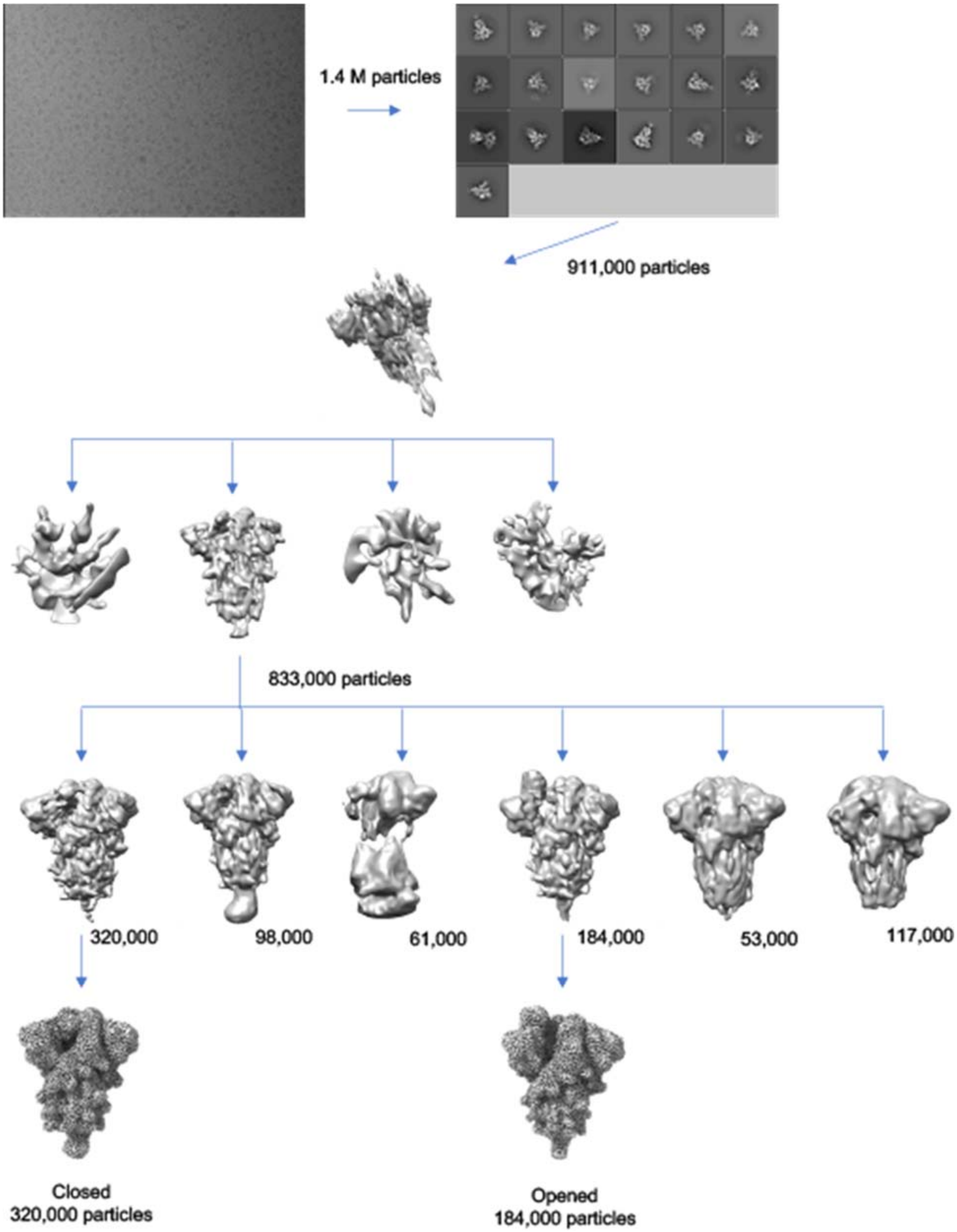


**Figure S7 Cryo-EM data processing workflow**

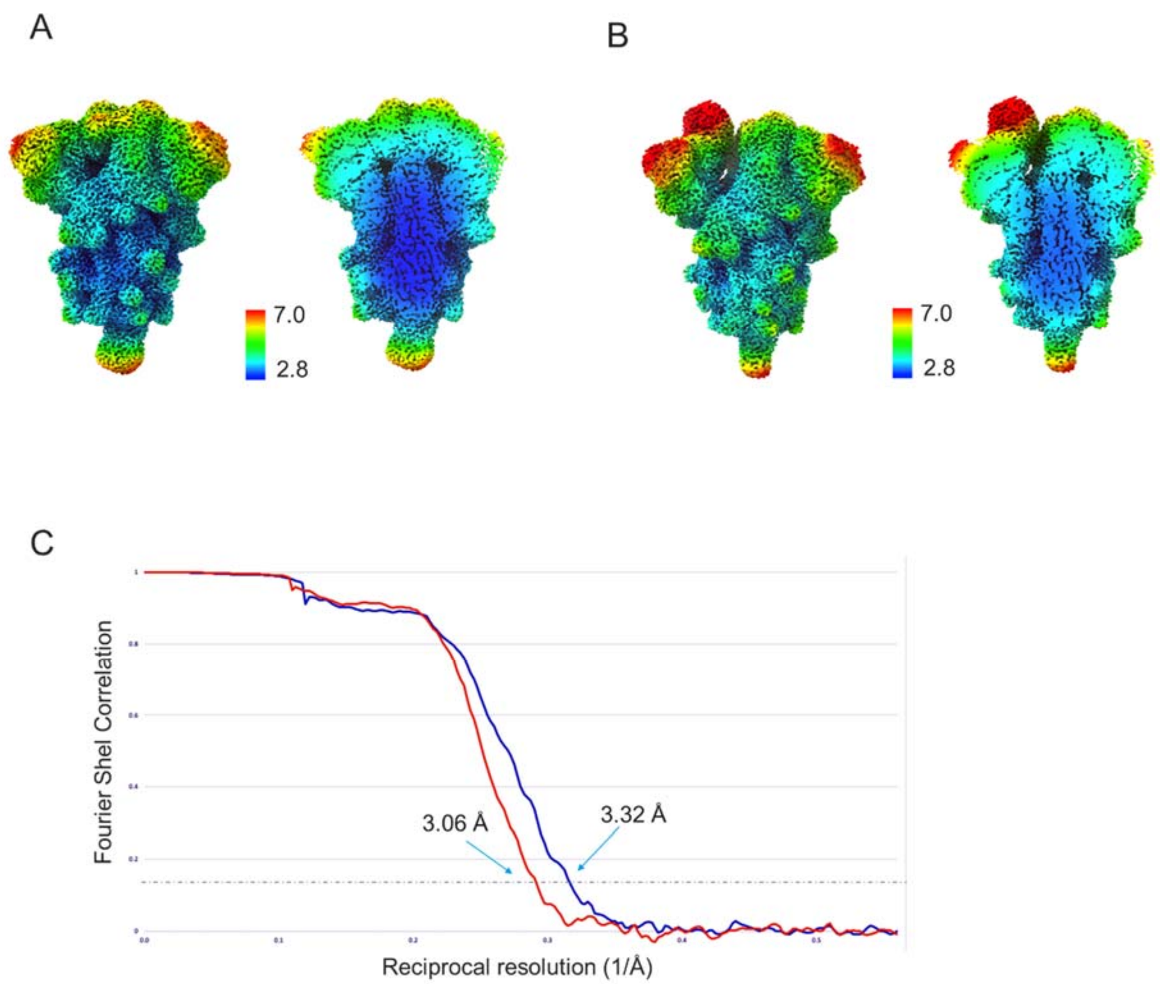


**Figure S8 Resolution assessment of cryo-EM structure**: A- Closed S-trimer map coloured by local resolution full map on the left and slice through the middle of the map on the right. B- Opened S-trimer map coloured by local resolution full map on the left and slice through the middle of the map on the right. C- Global resolution assessment by Fourier shell correlation at the 0.143 criterion.

**Supplementary Tables**

**Table S1. Thermal stabilities as measured with differential scanning calorimetry.**

| Constructs | Tm1 | Tm2 |
|---|---|---|
| ΔFurin | 48.91 | 63.97 |
| V987P | 48.27 | 64.08 |
| D614G | 48.57 | 65.95 |
| A944P | 48.52 | 64.02 |
| K986P V9787P (S-2P) | 48.33 | 64.38 |
| S-2P-D614N | 49.49 | 66.23 |
| S-2P-N532P | 48.7 | 64.53 |
| S-2P-T572I | 49.01 | 65.43 |
| S-2P-A892P | 48.96 | 65.01 |
| S-2P-DS1 | 51.76 | 64.61 |
| S-2P-A942P | 49.24 | 64.38 |
| A892P_A942P | - | 64.21 |
| S-closed+Fd | 55.02 | 66.32 |
| S-closed+Fd+K986P | 51.21 | 66.62 |
| S-closed | 51.65 | 66.31 |

**Table S2. SEC-MALS analysis**

| | Trimer peak | | | | LMW species | | | | Total |
|---|---|---|---|---|---|---|---|---|---|
| Mutation | Mw (kDa) | Rh (nm) | Mass fraction (%) | Retention time (min) | Mw (kDa) | Rh (nm) | Mass fraction (%) | Retention time (min) | Recovery (%) |
| S-closed | 465.6 | 9.165 | 89.8 | 5.319 | ND* | ND* | 8.7 | 5.647 | 96.7 |
| S-closed+Fd | 491 | 9.909 | 100 | 5.08 | - | - | - | - | 86.6 |

* Could not be determined due to low concentration.

**Table S3. Stability after freeze-thawing and SEC analysis**

| Mutation(s) | 0x FT | 1x FT | 3x FT | 5x FT |
|---|---|---|---|---|
| S-2P | 100 | 67 | 8 | 1 |
| S-2P+D614N | 100 | 83 | 43 | 12 |
| S-2P+DS1 | 100 | 75 | 24 | 4 |
| S-2P+N532P | 100 | 72 | 24 | 3 |
| S-2P+A892P | 100 | 80 | 21 | 3 |
| S-2P+A942P | 100 | 60 | 13 | 3 |
| S-2P+T572I | 100 | 81 | 22 | 3 |
| ΔFurin | 100 | 73 | 10 | 1 |
| D614G | 100 | 72 | 21 | 3 |
| A892P+A942P | 100 | 82 | 29 | 4 |
| S-closed+Fd+K986P | 100 | 77 | 31 | 11 |
| S-closed+Fd | 100 | 74 | 34 | 8 |
| S-closed | 100 | 78 | 55 | 35 |

#### Table S4 Cryo-EM data collection and refinement statistics

| | Closed trimer |
|---|---|
| **EMDB** | XXXX |
| **Magnification** | 105,000 |
| **Pixel size (Å)** | 0.86 |
| **Voltage (kV)** | 300 |
| **Electron exposure ($e^-$/Å$^2$)** | 65 |
| **Frames** | 50 |
| **Defocus Range (μm)** | -0.6 to -2.5 |
| **Number of movies collected** | 9,760 |
| **Particles extracted / Final** | 1,400,000 / 320,000 |
| **Symmetry** | N/A |
| **Masked resolution at FSC=0.143 (Å)** | 3.06 |
| **Map sharpening B-factor (Å$^2$)** | -65 |
| **PDB** | XXXX |
| **Model compositions** | |
| **Amino Acids** | 2916 |
| **Glycans** | N/A |
| **Mean B factors (Å2)** | |
| **Amino Acids** | 35 |
| **Glycans** | N/A |
| **R.m.s. deviations** | |
| **Bond lengths (Å)** | 0.004 |
| **Bond angles (°)** | 0.662 |
| **Validation** | |
| **MolProbity score** | 1.64 |
| **Clashscore** | 5.69 |
| **Rotamers outliers (%)** | 0.12 |
| **EMRinger score** | 2.82 |
| **Ramachandran plot** | |
| **Favored (%)** | 95.25 |
| **Allowed (%)** | 4.64 |
| **Outliers (%)** | 0.11 |
